# A conserved *Plasmodium* protein that localizes to liver stage nuclei is critical for late liver stage development

**DOI:** 10.1101/2022.12.13.519845

**Authors:** Debashree Goswami, Silvia A. Arredondo, William Betz, Janna Armstrong, Kenza M. Z. Oualim, Annette M. Seilie, Sean C. Murphy, Stefan H. I. Kappe, Ashley M. Vaughan

## Abstract

Malaria, the disease caused by *Plasmodium* parasites, causes significant mortality and morbidity. Whole parasite vaccination with pre-erythrocytic parasite stages, attenuated through sporozoite irradiation or chemo-attenuation, confers sterilizing immunity against subsequent parasite infection. This provides a rationale for the creation of whole parasite vaccines that are attenuated using gene editing. Here, we report on the creation of a novel genetically attenuated parasite (GAP) by the deletion of *Plasmodium LINUP,* encoding a <u>li</u>ver stage <u>nu</u>clear <u>p</u>rotein.

Epitope-tagging of LINUP in the rodent malaria parasite *Plasmodium yoelii* showed LINUP expression exclusively in liver stage nuclei after the onset of exo-erythrocytic schizogony. *P. yoelii* parasites with a gene deletion of *LINUP* (*linup* ^—^) suffered an exclusive liver stage phenotype with developmental arrested late in exo-erythrocytic schizogony. Liver stages showed incomplete segregation of nuclei and, mitochondria and apicoplast. These cellular perturbations caused a defect in exo-erythrocytic merozoite formation and a concomitant severe attenuation of liver stage-to-blood stage transition. *LINUP* gene deletion in *Plasmodium falciparum* also caused a severe defect in late liver stage differentiation. Importantly, *P. falciparum linup* ^—^ liver stages showed a severe defect in parasite transitioning from liver stage to viable blood stage infection. These results suggest that *P. falciparum LINUP*is a useful target for late liver stage attenuation and an additional gene deletion that can be incorporated into a late liver stage-arresting replication competent whole parasite vaccine.

## INTRODUCTION

### *Plasmodium* parasite species that infect humans are the causative agents of malaria

According to the World Health Organization’s latest World Malaria Report ^1^, there were an estimated 241 million malaria cases and 627,000 malaria deaths worldwide in 2020, mostly children in sub-Saharan Africa. Worryingly, this represents approximately 14 million more cases in 2020 as compared to 2019, and 69,000 more deaths.

RTS,S, a subunit vaccine that engenders antibody responses to the circumsporozoite protein (CSP) of *P. falciparum* (Pf), present on the pre-erythrocytic sporozoite stage, has been licensed for use in Africa. This partially protective vaccine has been reported to reduce malaria hospitalization by 21% and severe disease by 30% in African infants ^2^, but immunity wanes with time ^3^ and no data currently points to the prevention of new Pf infections or halting malaria transmission with RTS,S vaccination.

As an alternative to subunit vaccines, attenuated Pf whole parasite vaccines, that express thousands of potentially immunoprotective antigens, have proven efficacious in providing sterilizing protection from both controlled human malaria infection (CHMI) ^4–6^ and in endemic settings of malaria transmission ^7, 8^. Importantly, studies in rodent malaria models and nonhuman primate models show that attenuated parasite vaccinations engender not only neutralizing antibodies that prevent sporozoites from reaching the liver but also liver resident CD8^+^ T cells that eliminate liver stage-infected hepatocytes ^5, 9–21^. Thus, both arms of the adaptive immune response mediate parasite clearance at different points of infection, which results in superior protective immunity when compared to CSP antibody-based immunity engendered by current pre-erythrocytic subunit vaccines.

Attenuation is currently achieved in one of three ways – (i) irradiation of sporozoites that then arrest development into liver stages soon after hepatocyte invasion ^6, 22^, (ii) chemoprophylaxis that kills the liver stage parasite during liver stage development or after the liver stage-to-blood stage transition ^23, 24^ and (iii) genetic attenuation where genes essential for liver stage development are deleted from the parasite genome to create genetically attenuated parasites (GAP) ^10, 25, 26^. With respect to irradiated sporozoites, vialed, aseptic, irradiated, cryopreserved Pf NF54 strain sporozoites have been extensively used for intravenous vaccination and numerous successful clinical trials have been carried both in malaria-naïve individuals and in endemic regions of Africa^5–8, 27–29^. Wild type sporozoites have been used for vaccination in clinical trials in conjunction with chemoprophylaxis using pyrimethamine to kill developing liver stages early in their development, and chloroquine to kill the first wave of blood stage parasites that form after the emergence of exo-erythrocytic merozoites from the mature liver stage parasite ^24^.

GAPs are intrinsically attenuated and undergo complete liver stage arrest due to deletion of genes that are essential during liver stage development. This makes GAPs a feasible live-attenuated sporozoite vaccine candidate because it completely circumvents the need for chemoprophylaxis, thereby greatly reducing the complexity of clinical trial design. Furthermore, GAP can be designed to create liver stage parasites that arrest at any given time point during their development. GAPs were initially shown to be protective vaccines in rodent malaria models of infection using *P. berghei* (Pb) and *P. yoelii* (Py) GAP that arrested early in liver stage development ^30–32^. These first-generation rodent malaria GAP were early arresting replication deficient (EARD) and both antibodies and CD8^+^ T cells were shown to contribute to protection^33–39^. Knowledge of rodent malaria GAP design was used to create the first Pf NF54 strain human malaria EARD GAPS ^40^, with deletions in essential liver stage genes, including PfSPZ- GA1 (with deletions in *B9* and *SAP1*) and Pf GAP3KO, with deletions in *P52*, *P36* and *SAP1*. Pf GAP3KO was shown to be safe and completely attenuated in a pre-clinical trial ^41^. Furthermore, in a first-in-human clinical trial where Pf GAP3KO vaccination was administered by infectious mosquito bite, provided 50% sterile protection from a controlled human malaria infection (CHMI) by the administration of five infectious bites from mosquitoes harboring wildtype Pf NF54 ^25^. The additional EARD GAP candidate, PfSPZ-GA1, composed of aseptic, purified, cryopreserved sporozoites has also completed its first in-human clinical trials ^42^. PfSPZ-GA1 was administered by direct venous inoculation (DVI) and was shown to be completely attenuated. Intriguingly, vaccine efficacy of PfSPZ-GA1 against a five-mosquito bite CHMI with wildtype Pf NF54 was a modest 12% ^42^ and lower than that seen for Pf GAP3KO. Thus, Pf EARD GAP are safe, immunogenic and can provide sterile protection. Further studies on rodent malaria GAP have shown that GAP that are late liver stage-arresting replication competent (LARC) are more efficacious than EARD GAP and irradiated sporozoites in eliciting a protective immune response ^43, 44^, likely due to the increased antigen load and antigen breadth of the former. This observation led to the development of the first Pf NF54 LARC GAP which lacked the *PlasMei2* gene ^26^. *PlasMei2* GAP arrests late in liver stage development and does not transition from liver stage-to-blood stage in a human liver-chimeric model of injection, the FRG huHep humanized mouse ^26^. The equivalent rodent malaria Py GAP with deletion in *PlasMei2* is also severely attenuated but at very high immunization doses can transition from liver stage to blood stage in highly susceptible mice ^45^. Thus, it is important to identify additional genes for deletion that lead to late liver stage arrest of the parasite in order to ensure the safety of prospective LARC GAPs for use in humans.

Here we report on the identification of a second *Plasmodium* gene, *liver stage nuclear protein* (*LINUP*), that plays a conserved critical role in late liver stage development. Py *linup*^−^ parasites arrested late in liver stage development and showed severe attenuation in the transition to blood stage infection. Immunization of mice with Py *linup*^−^ sporozoites conferred complete protection from infectious sporozoite challenge. Pf *linup*^−^ parasites also arrested late in liver stage development and importantly, showed no viable liver-stage-to-blood stage transition in human-liver chimeric FRG huHep mice infused with human red blood cells. Thus, the *LINUP* gene deletion provides a second target for late liver stage attenuation and prevention of parasite transition from late live stage to blood stage infection and a such is useful for the refinement of replication competent parasite vaccines to achieve complete intrinsic attenuation.

## RESULTS

### Bioinformatic analysis of a human malaria late liver stage transcriptome for the discovery of genes expressed exclusively during liver stage development.

To identify novel genes that play critical roles only during liver stage development, we took advantage of a late liver stage *P. vivax* (Pv) transcriptome that was generated from infected livers of human-liver chimeric FRG huHep mice (manuscript in preparation). Mice were infected with one million Pv Thai field strain sporozoites isolated from *Anopheles dirus* mosquitoes. The mice were euthanized at day 8 after infection, late in liver stage development and the livers were processed for RNA extraction. Probes specific to the Pv exome were hybridized to purified RNA to enrich for parasite-specific transcripts, which were then converted to cDNA and bulk sequenced. Reads were normalized and expressed as reads <u>p</u>er <u>k</u>ilobase of transcript, per one <u>m</u>illion mapped reads (RPKM). This resulted in the capture of 4397 unique gene transcripts with RPKM values greater than two. To enrich for liver stage specific reads, the Pv RPKM reads were expressed as a fold change over the maximal RPKM reads for the orthologous Pf genes expressed during the blood stage of the life cycle (manuscript in preparation). This enabled us to compare and contrast asexual Pv liver stage development with asexual Pf blood stage development and specifically identify conserved transcripts that were highly expressed only during late liver stage development. The top ten genes from this analysis (**Table 1**) included *LISP1* ^46^, *LISP2* ^47–49^, *PALM* ^50^ and *SIAP2* ^51^, genes already known to be expressed during liver stage development and not during blood stage development. *Malonyl CoA-acyl carrier protein transacylase* was also detected, an enzyme involved in type II fatty acid biosynthesis, a pathway known to be expressed during Pf liver stage development and critical for Py and Pb liver stage development ^52–55^. We next applied a further criterion that genes had to have Py orthologs, which enabled us to use this rodent malaria model to assess liver stage phenotypes in mice with relative ease. In consequence, of the remaining five uncharacterized genes in the top ten, four candidates were prioritized for further analysis: a *protoporphyrinogen oxidase*, PY17X_0513300, and three hypothetical genes, PY17X_1003700, PY17X_1465200 and PY17X_1369800. PlasmoDB was then used to study the transcriptional profiles of the Py and Pf candidate orthologs across the life cycle as well as evidence of protein expression and gene essentiality in the Pf blood stage ^56^. This suggested that although PY17X_051300/PF3D7_102800 was dispensable in Pf asexual blood stages, the protein was expressed in the Pf blood stage gametocytes and the Py oocyst sporozoites and thus, the gene was not studied further. Analysis of PY17X_1003700/PF3D7_0404600 suggested the gene was dispensable in Pf asexual blood stages, but the protein was expressed in both the Pf blood stage merozoites and sporozoites and the gene was not studied further. Of note, analysis of PY17X_1369800/PF3D7_1351300 revealed no evidence of protein expression in either Py or Pf datasets. Additionally, analysis of PY17X_1465200/PF3D7_1249700 suggested the gene was dispensable for Pf asexual blood stage replication and expressed in the Py liver stage at 40 hours of development. Therefore, PY17X_1369800/PF3D7_1351300 and PY17X_1465200/PF3D7_1249700 underwent further study. Gene knockout of PY17X_1369800 was achieved but the phenotype of the knockout was equivalent to wildtype across the life cycle and thus this gene deletion was not pursued further (data not shown).

**TABLE 1.** Top ten Plasmodium vivax genes expressed during day eight of liver stage development based on a comparison with Plasmodium falciparum asexual blood stages

| *Plasmodium vivax*<br>gene identifier | Gene product | *Plasmodium vivax* day 8<br>liver stage<br>RPKM | RPKM<br>*Plasmodium vivax* fold<br>change<br>compared to<br>mean RPKM<br>for<br>*Plasmodium falciparum*<br>asexual<br>blood stage | *Plasmodium falciparum*<br>ortholog | *Plasmodium yoelii* ortholog |
| --- | --- | --- | --- | --- | --- |
| PVX_122435 | malonyl CoA-acyl<br>carrier protein<br>transacylase | 857 | 143 | PF3D7_1312000 | PY17X_1412300 |
| PVX_001010 | conserved<br>hypothetical protein | 1489 | 68 | PF3D7_0404600 | PY17X_1003700 |
| PVX_000975 | liver specific<br>protein 2 (LISP2) | 653 | 61 | PF3D7_0405300 | PY17X_1004400 |
| PVX_085550 | liver specific<br>protein 1 (LISP1) | 717 | 54 | PF3D7_1418100 | PY17X_1027000 |
| PVX_101370 | conserved<br>hypothetical protein | 375 | 53 | PF3D7_1249700 | PY17X_1465200 |
| PVX_083460 | conserved<br>hypothetical protein | 86 | 42 | PF3D7_1351300 | PY17X_1369800 |
| PVX_088860 | sporozoite<br>invasion-<br>associated protein<br>2 (SIAP2) | 259 | 37 | PF3D7_0830300 | no ortholog |
| PVX_111315 | protoporphyrinogen<br>oxidase | 110 | 29 | PF3D7_1028100 | PY17X_0513300 |
| PVX_086345 | conserved<br>hypothetical protein | 71 | 28 | PF3D7_1401700 | no ortholog |
| PVX_113280 | liver merozoite<br>formation protein<br>(PALM) | 103 | 25 | PF3D7_0602300 | PY17X_0102700 |

### Py LINUP is expressed exclusively during liver stage development

The remaining candidate, PY17X_1465200, is a single exon gene encoding a 746 amino acid protein and is conserved among *Plasmodium* species (**Figure 1A**). The overall amino acid identity between the Py, Pf and Pv syntenic orthologs was 40%, whilst amino acid similarity was 60%. Identity in a 122 amino acid stretch near to the N-terminus (amino acids 44-161) was 89%. In addition, comprehensive protein BLAST searches revealed that the gene has no orthologs in other *Apicomplexa* or any other eukaryote and is thus unique to *Plasmodium*. Further analysis of the PY17X_1465200 amino acid sequence revealed a conserved N-terminal 24 amino acid nuclear localization sequence (NLS) in Py, Pf and Pv that was nearly identical in amino acid sequence among the three species (**Figure 1A**). Based on the presence of the NLS and the liver stage nuclear localization of PY17X_1465200 (see below), we named the gene *<u>li</u>ver stage <u>n</u>uclear <u>p</u>rotein* (*LINUP*). To determine the spatial-temporal expression of Py LINUP, a transgenic parasite line was created with an mCherry tag fused to the C-terminus of the endogenous Py *LINUP* (**Figure 1B**, top panel cartoon). Standard transfection procedures ^57^ resulted in successful double crossover homologous recombination and transgenic parasites were cloned. Integration of the mCherry tag was confirmed by transgene-specific PCR (**Figure S1A, B**). Py *LINUP* mCherry-tagged parasite clones (Py LINUP^mCherry^) were comparable to wildtype during mosquito stage development and produced comparable oocyst numbers, oocyst prevalence and salivary gland sporozoite numbers per mosquito (**Figure S1C**). In addition, the liver stage development and the transition to blood stage patency was not affected by the mCherry tag since the intravenous injection of 10,000 salivary gland sporozoites from both Py wildtype and Py LINUP^mCherry^ parasites into groups of Swiss Webster mice, resulted in all mice becoming blood stage patent on the third day after sporozoite injection (**Figure S1D**).

**Figure 1:**
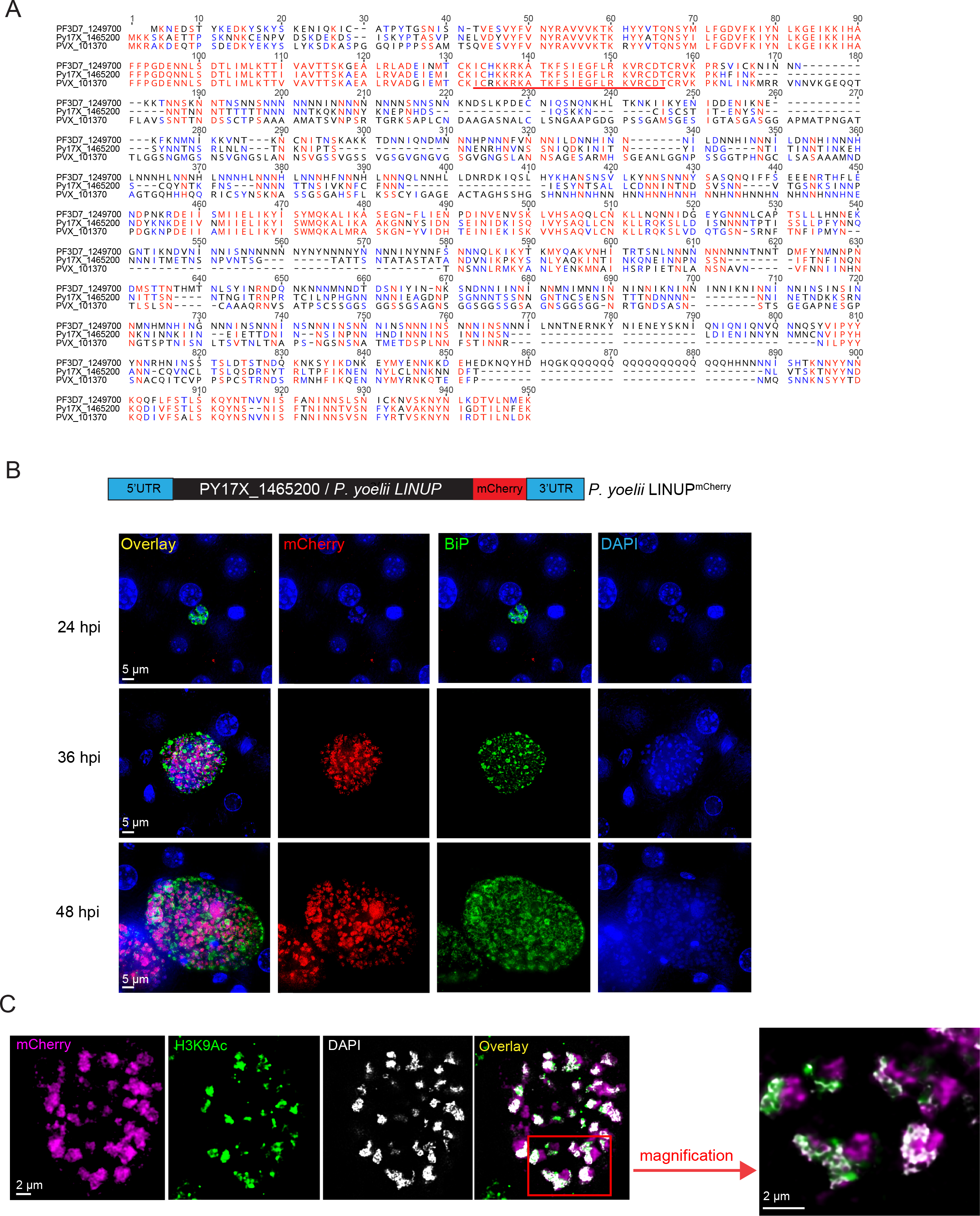
Py <u>Li</u>ver stage <u>Nu</u>clear <u>P</u>rotein (LINUP) (PY17X_1465200) localizes to the **nucleus of liver stage parasites.** A. Amino acid alignment between Pf LINUP (PF3D7_1249700) and its syntenic orthologs Py LINUP (PY17X_1465200) and *P. vivax* LINUP (PVP01_1466800) shows 40% sequence identity and 60% sequence similarity. 80 – 100% sequence similarity is shown in red and 60 – 80% sequence similarity is shown in blue. The predicted and conserved nuclear localization sequence (NLS) is underlined in red and spans amino acids 131-154. B. To visualize the localization of Py LINUP, an mCherry epitope-tagged parasite strain, Py LINUP^mCherry^ was generated by fusing an mCherry tag to the LINUP C-terminus. The tagged transgenic parasite replaces the endogenous *LINUP* with the tagged copy. Immunofluorescence assays (IFAs) on Py LINUP^mCherry^ infected mouse livers using an mCherry antibody, an antibody to the endoplasmic reticulum marker BiP and DAPI to delineate DNA indicate that the protein is expressed after 24 hours of infection and at both 36 and 48 hours localizes to the nucleus of liver stage schizonts. Scale bar size is 5 μm. C. At 48 hours, LINUP shows partial co-localization with the acetylated histone H3 marker lysine 9 and DAPI, scale bar is 5 μm, which is further indicated in the magnified image to the right where the scale bar size is 2 μm.

Expression of LINUP^mCherry^ was monitored across the complete life cycle by immunofluorescence assay (IFA). Interestingly, LINUP^mCherry^ expression was only seen during mid-to-late liver stage development, which in Py has an approximate 50-hour duration.

LINUP^mCherry^ expression was not detected at 24 hours of liver stage development (**Figure 1B**) but was detected at both 36 and 48 hours (**Figure 1B**), the latter a timepoint when merozoites form during the final stages of exo-erythrocytic schizogony. LINUP^mCherry^ localized to live stage nuclei together with parasite DNA and partially co-localized with the histone marker, histone 3 acetylated lysine 9 (H3K9) which marks areas of active gene expression (**Figure 1C**).

### Py *LINUP* plays an important role in liver stage development

To study the importance of LINUP function, we used CRISPR/Cas9 technology to delete the *LINUP* gene from the Py genome (**Figure 2A**). After blood stage schizont transfection, *in vivo* positive drug selection with pyrimethamine in mice and isolation of drug resistant parasites, the gene knockout was confirmed by PCR on isolated genomic DNA, using primer pairs diagnostic for the gene deletion. Py *linup*^−^ parasites were then cloned, and two clones, c3 and c5, were used for phenotypic analysis (**Figure 2B**) across the parasite life cycle. The two clones were comparable to wildtype parasites with respect to asexual blood stage growth (**Figure 2C**). The development of mosquito stages of the life cycle was also comparable based on enumeration of oocyst numbers (**Figure 2D**), oocyst prevalence (**Figure 2E**) and salivary gland sporozoite numbers (**Figure 2F**).

**Figure 2:**
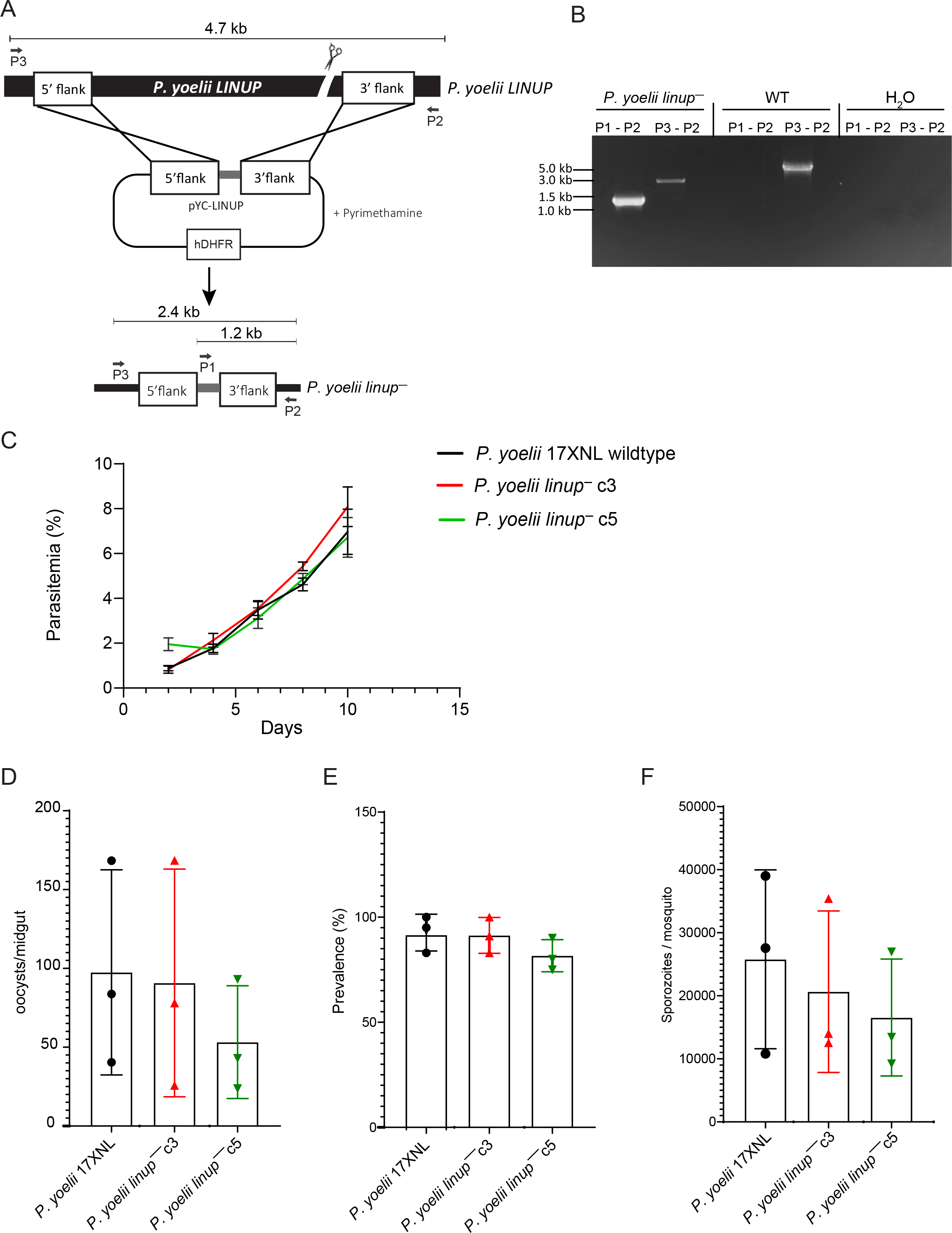
Py *linup^−^* creation and analysis of mosquito stage development. A. The schematic depicts the generation of the Py *linup^−^* parasite using CRISPR/Cas9-mediated gene editing. **B.** Agarose gel electrophoresis shows the PCR products corresponding to the gene deletion of Py LINUP in clone 3. Primers used to verify the gene deletion are indicated and the sizes of the PCR amplicons are shown in kilobases. The same result was seen for clone 5. C. Blood stage growth was compared in groups of five Swiss Webster mice for Py XNL wildtype and Py *linup^−^* clones c3 and c5. One million infected red blood cells were injected intravenously into each mouse and parasitemia measured every other day for ten days. Growth rates were comparable suggesting LINUP has no essential role in blood stage replication. Data is represented as mean +/- SEM, *n*=5 biological replicates. Statistical analysis was carried out using two-way ANOVA using Tukey’s multiple comparison test. P>0.05 is taken as not significant. Py *linup^−^* clones c3 and c5 did not have defects in mosquito infectivity as counts for D., oocysts/midgut, E., oocyst prevalence and F., salivary gland sporozoites/mosquito were comparable between Py *linup^−^* c3 (red) and c5 (green) to Py wildtype (black). Data is represented as mean +/- SD, *n*=3 biological replicates. Statistical analysis was carried out using two-way ANOVA using Tukey’s multiple comparison test. P>0.05 is taken as not significant.

To determine if liver stage development was affected by the deletion of *LINUP*, Py wildtype and *linup*^−^ sporozoites were isolated from infected *Anopheles stephensi* mosquito salivary glands and injected intravenously into groups of BALB/cJ and BALB/cByJ mice. These mouse strains are both susceptible to Py infection with BALB/cByJ mice being more susceptible than BALB/cJ mice ^58^. The time to blood stage patency, in days, was then determined from Giemsa-stained thin blood smears. A dose escalation study in BALB/cJ using 1,000, 10,000 or 50,000 sporozoites per dose, revealed evidence of severe liver stage attenuation. Wildtype sporozoite-infected mice all became patent either on day four (1,000 sporozoites), days three-four (10,000 sporozoites) or day three (50,000 sporozoites) post infection. In contrast, in Py *linup*^−^ sporozoite infection with 1,000 sporozoites only one of ten mice became patent and this with delay (day twelve) and seven of twenty mice became patent when infection was done with 10,000 sporozoites with patency in the blood stage positive mice delayed to days seven through twelve. Five of ten mice became patent when using the 50,000 sporozoite dose and patency was delayed to days seven to nine (**Table 2**). Highly susceptible BALB/cByJ mice infected with 50,000 wildtype sporozoites all became patent on day three after sporozoite infection, whereas only nine of twenty mice infected with Py *linup*^−^ sporozoites became patent and the day to patency ranged from days eight to ten (**Table 2**). Thus, Py *linup*^−^ parasites frequently could not establish successful blood stage infection. In those mice that did become blood stage patent, we estimated that there was a greater than 99% reduction in Py *linup*^−^ pre-erythrocytic infection, using assumptions published previously ^59, 60^.

**TABLE 2.** Attenuation of P. yoelii linup^−^ liver stage development in BALB/c mice

|  |  |  |  |  |
| --- | --- | --- | --- | --- |
| Parasite genotype | Mouse strain | Number of sporozoites<sup>b</sup> | Number of mice patent/number of mice infected<sup>c</sup> | Days to patency (number of mice)<sup>d</sup> |
| *P. yoelii* 17XNL | BALB/cJ | 1,000 | 5/5 | 4 |
|  |  | 10,000 | 5/5 | 3 (2) |
|  |  |  |  | 4 (3) |
|  |  | 50,000 | 3/3 | 3 |
|  | BALB/cByJ | 50,000 | 3/3 | 3 |
| *P. yoelii linup*<sup>-a</sup> | BALB/cJ | 1,000 | 1/10 | 12 |
|  |  | 10,000 | 7/20 | 7 (1) |
|  |  |  |  | 9 (2) |
|  |  |  |  | 10 (1) |
|  |  |  |  | 12 (3) |
|  | BALB/cJ | 50,000 | 5/10 | 7 (2) |
|  |  |  |  | 8 (1) |
|  |  |  |  | 9 (2) |
|  | BALB/cByJ | 50,000 | 9/20 | 8 (3) |
|  |  |  |  | 9 (2) |
|  |  |  |  | 10 (4) |

<sup>a</sup>Clone c3 of *Py linup*<sup>-</sup> was used for analysis.
<sup>b</sup>Salivary gland sporozoites were isolated from infected *Anopheles stephensi* mosquitoes, and mice were challenged intravenously with the listed number of sporozoites.
<sup>c</sup>The number of patent mice per number of mice challenged is indicated. Detection of blood stage patent parasitemia was carried out by Giemsa-stained thin blood smear. Attenuation was considered complete if mice remained blood stage negative for 21 days.
<sup>d</sup>If mice became blood stage patent, the day to patency is listed, with the number of mice that became patent in parentheses.

### Py *linup*^−^ parasites arrest late in liver stage development

To further study the defects of Py *linup*^−^ parasites in the pre-erythrocytic stages of infection, BALB/cByJ mice were infected intravenously with 250,000 Py *linup*^−^ sporozoites. Mice were euthanized at different time points after infection (24, 36 and 48 hours), livers were removed, perfused, fixed, sectioned, and infected tissue sections subjected to IFA using parasite-specific antibodies (**Figure 3**). Liver stage size measurements revealed that at 24 hours post infection, Py *linup*^−^ liver stages were comparable in size to wildtype (**Figure 3A**).

**Figure 3:**
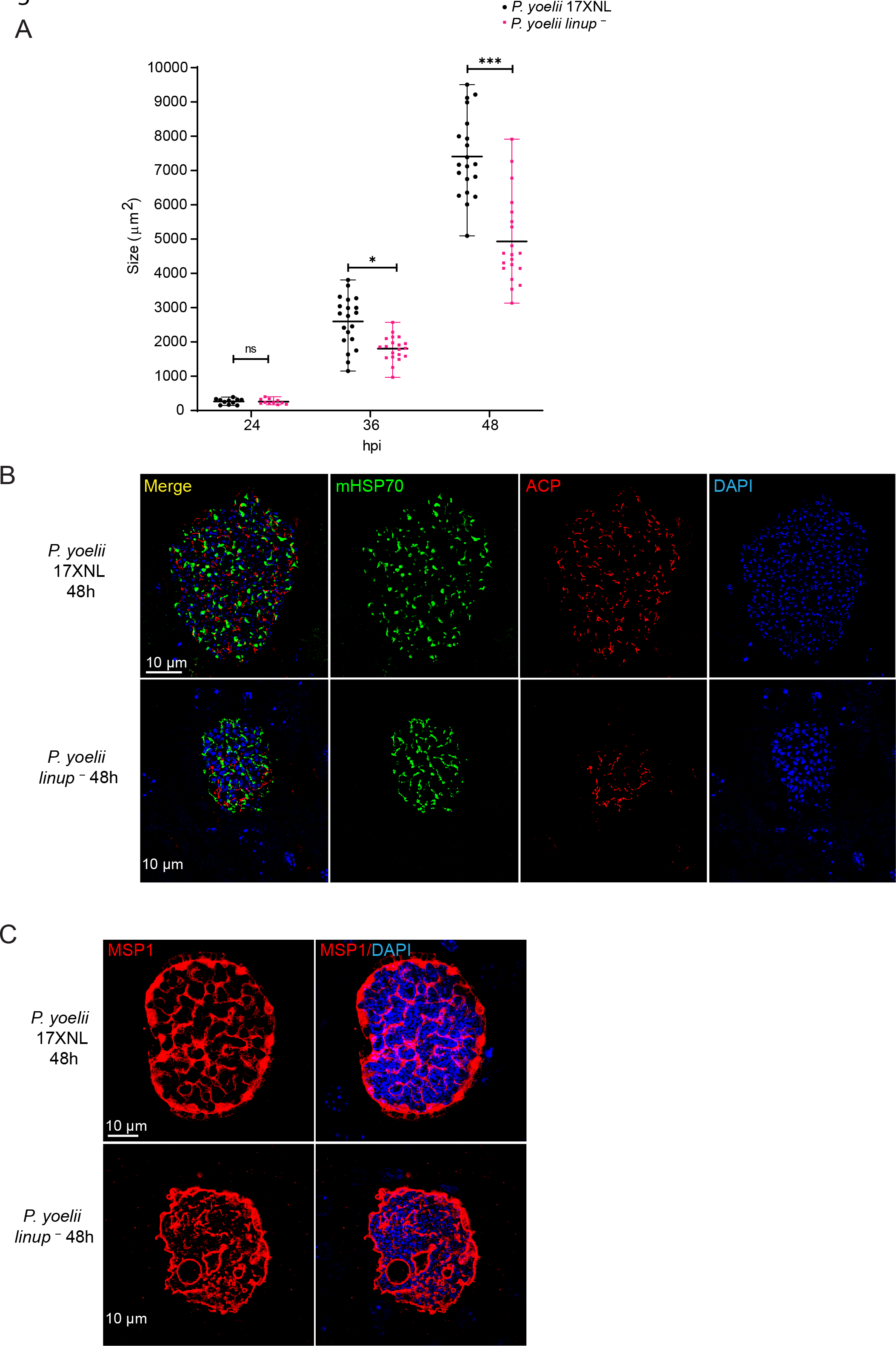
Analysis of Py *linup^−^* liver stage development. Tissue sections were prepared from BALB/cJ mice infected with 250,000 sporozoites of either Py wildtype or Py *linup^−^* at 24, 36 and 48 hours post infection and analyzed by IFA. A. Comparison of the size of liver stage parasites (based on area at the parasite’s largest circumference) between Py wildtype and Py *linup^−^* at 24, 36 and 48 hours indicate that Py wildtype liver stage schizonts are significantly larger than Py *linup^−^* at 36 and 48 hours. Data is represented as mean ± SD. Each datapoint refers to the mean size of at least 20 parasites for each timepoint. Statistical analysis was carried out using two-way ANOVA using Tukey’s multiple comparison test. * = P.05, ***P<0.001, P>0.05 is taken as ns. Liver stage development was compared at 48 hours using antibodies against B., the parasite mitochondria (mHSP70, green) and apicoplast (ACP, red), and B., the parasite plasma membrane and mature exo- erythrocytic merozoite marker, MSP1 DNA was stained with DAPI (blue). Scale bar: 10 μm. Py *linup^−^* liver stage parasites are smaller compared to Py wildtype and display less branching of the mitochondria and apicoplast organelles. Py *linup^−^* expresses the late liver stage proteins MSP1, however there is aberrant segregation of cytomeres and incomplete formation of mature exo-erythrocytic merozoites.

However, at both 36 and 48 hours post infection, Py *linup*^−^ liver stages were significantly smaller than wildtype with the size differences being more pronounced at the later time point post infection (**Figure 3A**). At 48 hours post infection, differences in protein expression and DNA replication/segregation were evident in Py *linup*^−^ liver stages when compared to wildtype liver stages. Specifically, the branching of the mitochondria and apicoplast was reduced and DNA replication as well as DNA segregation appeared significantly reduced (**Figure 3B**).

Furthermore, the extensive invaginations of the liver stage plasma membrane that precedes merozoite formation called cytomere formation was severely disordered in Py *linup*^−^ liver stages. This was evident from the changed expression pattern of the parasite plasma membrane marker MSP1, which localizes to the late liver stage schizont plasma membrane (**Figure 3C**).

### Immunization with Py *linup*^−^ provides protection from sporozoite challenge

To determine if vaccination of BALB/cJ mice with Py *linup*^−^ sporozoites protects from infectious sporozoite challenge, mice were intravenously boosted 33 days after first immunization with either 1,000 or 10,000 Py *linup*^−^ sporozoites or mock immunized with an equivalent volume of salivary gland extract from uninfected mosquitoes (**Table 3**). Only mice that did not become blood stage patent after Py *linup*^−^ sporozoite infection in studies of breakthrough (Table 2) described in the preceding section were used. After a further 34 days immunized mice were challenged with an intravenous injection of 10,000 wildtype sporozoites. Mice immunized with salivary gland extract all became blood stage patent three to four days after challenge (**Table 3**). Six of nine mice immunized twice with 1,000 Py *linup*^−^ sporozoites were protected from challenge and the remaining three mice showed a significant delay to onset of blood stage patency, becoming patent on days eight to ten after challenge (**Table 3**). All mice (thirteen) immunized twice with 10,000 Py *linup*^−^ sporozoites were protected after challenge.

**TABLE 3.** *P. yoelii linup^−^* immunization protects from a wildtype sporozoite challenge

|  | Number of sporozoites used for immunization/challenge |  |  |  |
| --- | --- | --- | --- | --- |
| Parasite genotype | Prime<sup>a</sup> | Boost (days after prime)<sup>a</sup> | Challenge (days after boost)<sup>b</sup> | Number of mice protected (days to patency)<sup>c</sup> |
| N/A | salivary gland extract | salivary gland extract | 10,000 | 0/5 (3, 4) |
| *P. yoelii linup<sup>-</sup>* | 1,000 | 1,000 (33) | 10,000 (34) | 6/9 (8,10) |
| *P. yoelii linup<sup>-</sup>* | 10,000 | 10,000 (33) | 10,000 (34) | 13/13 |

<sup>a</sup>*P. yoelii linup<sup>-</sup>* salivary gland sporozoites were isolated from infected *Anopheles stephensi* mosquitoes, and BALB/cJ mice were immunized intravenously with the listed number of sporozoites. The days after the prime that the boosts took place is indicated in parentheses.
<sup>b</sup>Mice were challenged intravenously with wildtype salivary gland sporozoites. The days after the boost the challenge took place are indicated in parentheses.
<sup>c</sup>The number of protected mice per number of mice challenged is indicated and the days to patency are indicated in parentheses. Protection was considered complete if mice remained blood stage negative for 21 days after challenge, based on Giemsa-stained thin blood smear.

This result shows that immunization with Py *linup*^−^ GAP engenders an immune response that protects mice from a wildtype sporozoite challenge.

### Pf *linup*^−^ parasites show severe attenuation late in liver stage development

We next created a Pf *linup^−^* parasite line using a CRISPR/Cas9 methodology for gene deletion (**Figure 4A**) ^26^. After plasmid transfection into the parental strain Pf NF54, and positive selection, gene knockout parasites were confirmed by PCR and subsequently cloned (**Figure 4B**). Two individual clones, B1 and B4, were isolated for phenotypic characterization across the life cycle using the parental wildtype NF54 as a comparator. Asexual blood stage replication and gametocytogenesis of the Pf *linup^−^* clones B2 and B4 were comparable to wildtype Pf NF54 parasites (data not shown). Similarly, mosquito stage development for the two clones based on oocyst counts (**Figure 4C**), oocyst prevalence (**Figure 4D**), and ultimately the number of salivary gland sporozoites per mosquito (**Figure 4E**), was comparable to wildtype Pf NF54. Subsequently, one million salivary gland sporozoites were injected into human hepatocyte-chimeric FRG NOD huHep mice ^61^, to analyze Pf *linup^−^* liver stage burden at day seven (mature- liver stage) by qPCR and liver stage development at days five (mid-liver stage) and seven by IFA of infected liver tissue (**Figure 5**). Parasite liver stage burden appeared higher in wildtype when compared to Pf *linup^−^* on day seven, based on qRT-PCR of Pf 18S rRNA (**Figure 5A)**.

**Figure 4:**
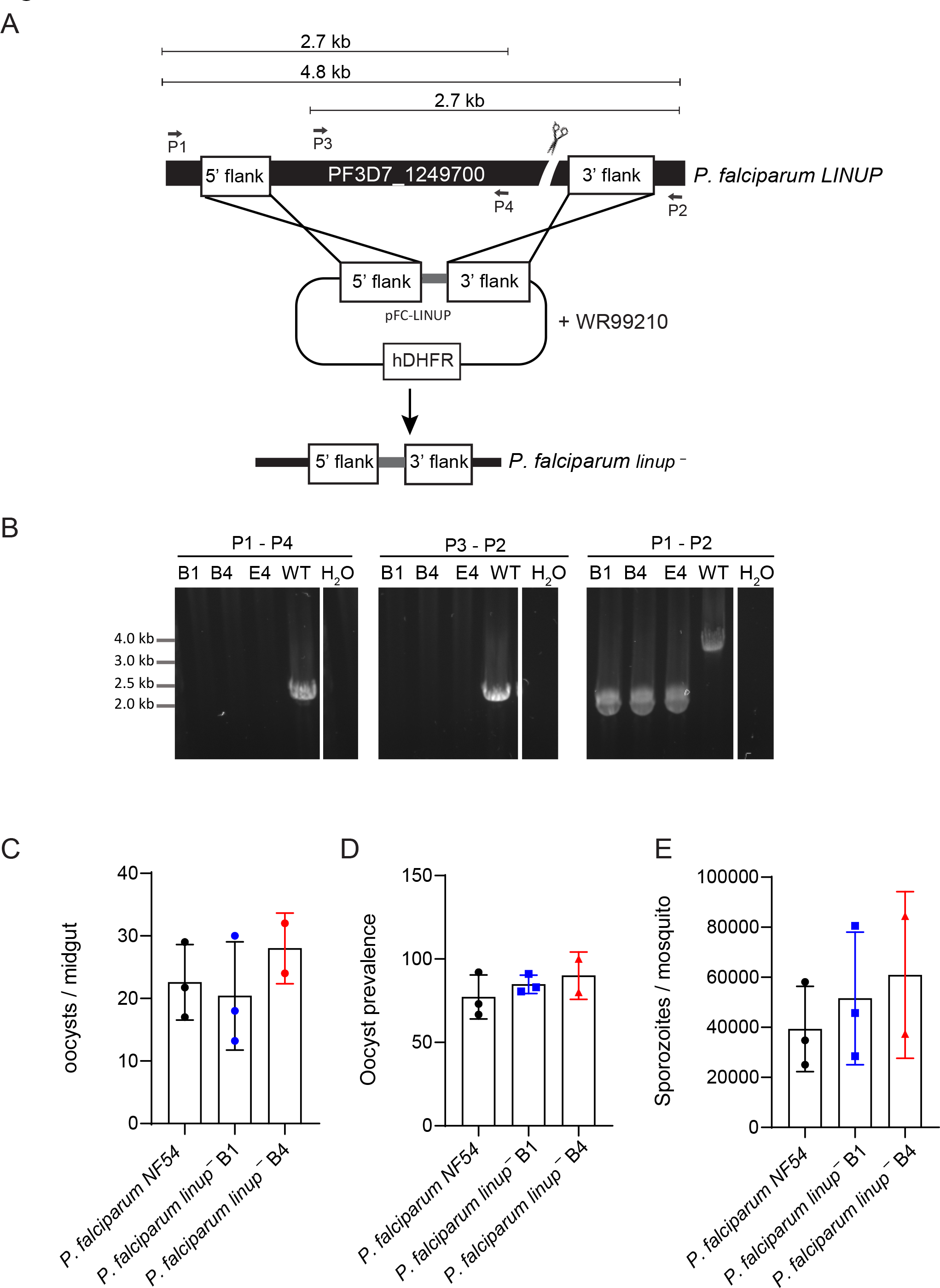
Pf *linup^−^* creation and analysis of mosquito stage development. A. The schematic depicts the generation of Pf *linup^−^* knockout parasites using CRISPR/Cas9-mediated gene editing. Primers used to verify the gene deletion are indicated and the sizes of the PCR amplicons are shown in kilobases. B. Agarose gel electrophoresis shows the PCR products corresponding to the gene deletion of Pf LINUP in clones B1 and B4. Pf *linup^−^* did not have defects in mosquito infectivity as counts for C., oocysts/midgut, D., oocyst prevalence and E., salivary gland sporozoites/mosquito were comparable between Pf *linup^−^* clones B1 (blue) and B4 (red) and Pf NF54 (black). Data is represented as mean +/- SD, *n*=2 biological replicates. Statistical analysis was carried out using two-way ANOVA using Tukey’s multiple comparison test. P>0.05 is taken as not significant.

**Figure 5:**
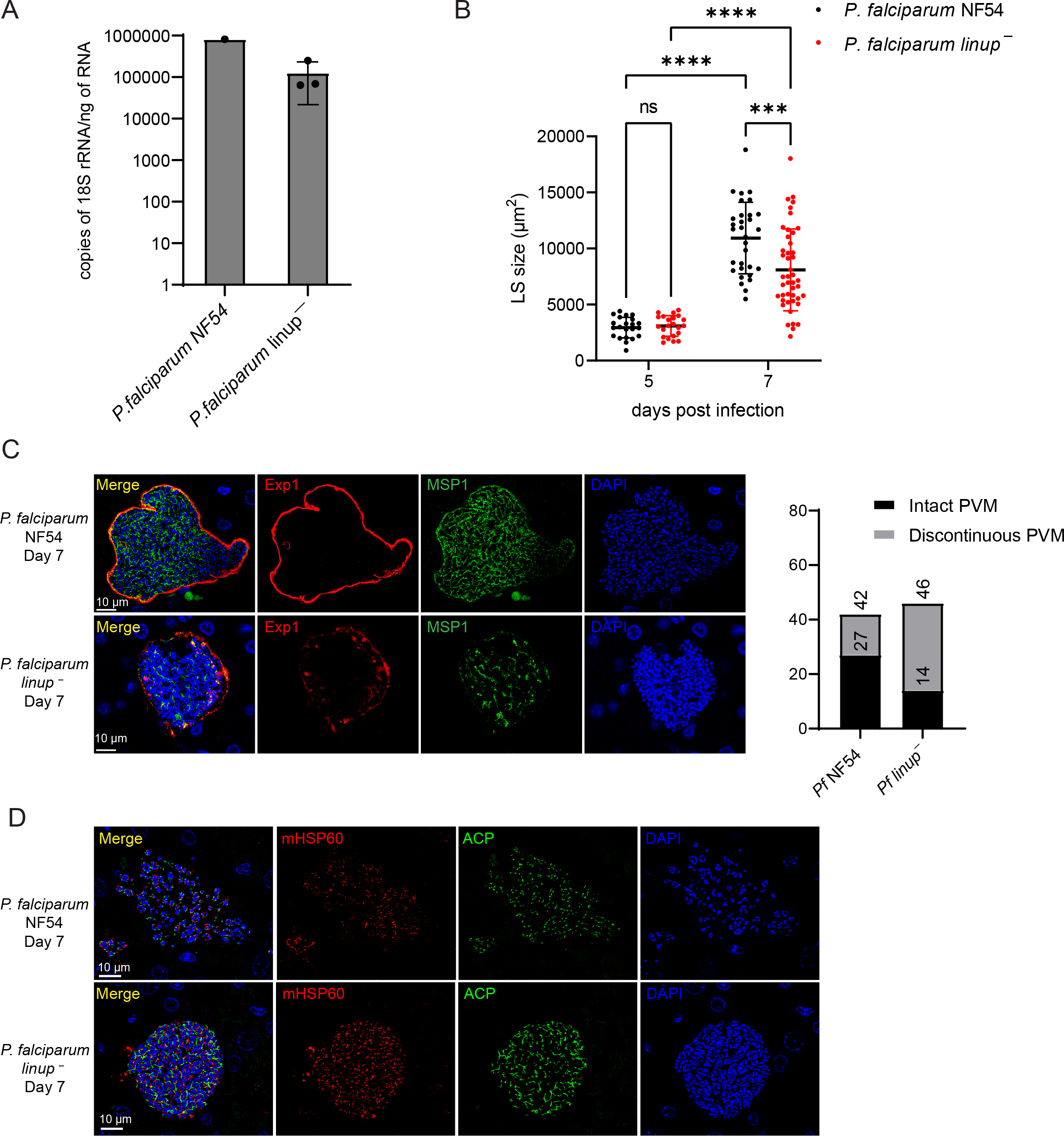
**Pf *linup^−^* liver stages display normal growth but aberrant late liver stage development**. A. One million sporozoites from Pf NF54 and Pf *linup^−^* were injected into one and three FRG NOD huHep mice, respectively and livers were harvested on day seven. Parasite liver load was quantified using 18S rRNA qRT-PCR on extracted RNA from infected livers. Data for Pf *linup^−^* injected mice are represented as mean ± SD. B. Tissue sections were prepared from FRG NOD huHep mice that were infected with one million sporozoites of Pf NF54 and Pf *linup^−^* on days 5 and 7 post-sporozoite infection and analyzed for size and protein expression. B. Comparison of the size of liver stage parasites (based on area at the parasite’s largest circumference) between Pf NF54 and Pf *linup^−^* on days 5 and 7 post-sporozoite infection. No growth defect was seen in Pf *linup^−^* mid- liver stage schizonts on day five, however a significant growth defect was observed in late-liver stage schizonts on day seven. Data is represented as mean ± SD. Each datapoint refers to the mean size of at least 30 parasites for each timepoint. Statistical analysis was carried out using two-way ANOVA using Tukey’s multiple comparison test. ***P<0.001, P>0.05 is not significant (ns). C . Left panel: IFA shows that at seven days of infection the wildtype NF54 parasitophorous vacuole membrane (PVM) protein EXP1 had a circumferential and contiguous staining pattern in the wildtype, as expected. In comparison EXP1 expression in mature Pf *linup^−^* day seven liver stages was patchy and aberrant. Right Panel: Quantification of late liver stage schizonts with regard to a contiguous or discontinuous PVM based on the localization of the PVM marker EXP1 showed 64% (27 out of 42) wildtype liver stages displayed a contiguous PVM while only 30% (14 out of 46) Pf *linup^−^* displayed a contiguous PVM. D. IFA shows that at seven days of infection the parasite mitochondria protein HSP60 (mHSP60) and the apicoplast acyl carrier protein (ACP) in wildtype NF54 undergoes segregation in exo-erythrocytic stage merozoites, while these organelles remain branched in the Pf *linup^−^* liver stage parasite.

We thus analyzed liver stage size during development. Deletion of Pf LINUP did not affect the size of mid-liver stage schizonts since the size of Pf *linup^−^* liver stages were comparable to wildtype on day five **(Figure 5B)**. However, deletion on Pf LINUP resulted in a significant growth defect during late liver stage schizogony since Pf *linup^−^* liver stages were significantly smaller on day seven, the time point of liver stage maturation (**Figure 5B)**. However, the ranges of liver stage size observed for Pf wildtype and Pf *linup^−^* liver stages overlapped significantly. To further investigate mature liver stage cellular differentiation, we employed IFAs to analyze development of the parasitophorous vacuole membrane (PVM) using an antibody to Pf EXP1, as well as development of the parasite plasma membrane (PPM), using an antibody to MSP1 (**Figure 5C**). Mature day seven wildtype liver stages showed circumferential EXP1 expression with a contiguous distribution on the PVM. MSP1 expression patterns were consistent with the complex invaginations of the plasma membrane during late liver stage cytomere formation. In contrast, EXP1 expression in Pf *linup^−^* day seven liver stages exhibited a patchy and disrupted pattern (**Figure 5C**). While 64% of wildtype liver stages on day seven displayed a contiguous circumferential distribution of EXP1 on the PVM, only 30% of Pf *linup^−^* displayed this phenotype (**Figure 5C**). Furthermore, MSP1 expression patterns indicated compromised cytomere formation (**Figure 5C**). Late liver stage schizogony involves the segregation of parasite organelles into individual exo-erythrocytic merozoites. To visualize the differentiation of parasite mitochondria and apicoplast, IFAs were performed using antibodies to mitochondrial HSP60 (mHSP60) and apicoplast-targeted acyl carrier protein (ACP). Wildtype late exo-erythrocytic schizonts on day seven displayed a discrete dotted localization of mHSP70 and ACP around nuclear centers, indicating segregation of these organelles into exo-erythrocytic merozoites (**Figure 5D**). In contrast, these organelles remained branched and tubular in the Pf *linup^−^* day seven liver stages (**Figure 5D**). These analyses show that LINUP is critical for late Pf liver stage exo-erythrocytic merozoite differentiation.

### Pf *linup*^−^ liver stages fail to transition to viable blood stage infection

We further assessed Pf *linup*^−^ liver stage attenuation and the impact of the observed liver stage phenotypes on parasite transition to blood stage infection ^26^. FRG NOD huHep mice were intravenously infected with one million Pf NF54 wildtype or one million *linup*^−^ sporozoites (**Figure 6A**). On days six and seven after sporozoite infection, mice were injected intravenously with human red blood cells (**Figure 6A**). The injection of human red blood cells enables the capture of viable invasive exo-erythrocytic merozoites released from mature liver stages and allows for the initiation of asexual blood stage replication. Parasite biomass after transition was first assessed by qRT-PCR measurement of 18S rRNA levels from the blood of mice exsanguinated seven days after sporozoite injection (**Figure 6B**). This assay has a reliable limit of detection above 20 parasites per ml of blood ^62^. At day seven, blood from the mouse injected with wildtype Pf NF54 sporozoites contained 4.8 × 10^6^ parasites per ml (**Figure 6B**). Mice injected with Pf *linup^−^* sporozoites ranged from 0 to 9.6 × 10^3^ parasites per ml, suggesting a severe liver stage attenuation. Since it has been hypothesized that viable parasites or nonviable parasite material released into the blood can be the source of 18S rRNA RNA into the blood ^26^, blood from the exsanguinated animals was placed into *in vitro* culture and sampled again seven days later (day 14 after sporozoite injection). Pf NF54 wildtype parasite replication during this time was significant, giving rise to 680 × 10^6^ parasites per ml of blood, a greater then 100-fold increase (**Figure 6B**). In addition, parasites were detected by Giemsa-stained thin blood smear. In contrast, the *in vitro* culture of blood from mice injected with Pf *linup^−^* sporozoites showed only small increases (the culture with no signal remained negative) (**Figure 6B**) and parasites were not detected by Giemsa-stained thin blood smear. This result suggests that the initial detection of parasite biomass in the blood from mice injected with Pf *linup^−^* sporozoites was due to the release of parasite rRNA from non-viable liver stage parasites into the blood stream of the mice and not viable exo-erythrocytic merozoites. Thus, the Pf *linup^−^* parasite is severely attenuated in the humanized mouse model of liver stage-to-blood stage transition.

**Figure 6:**
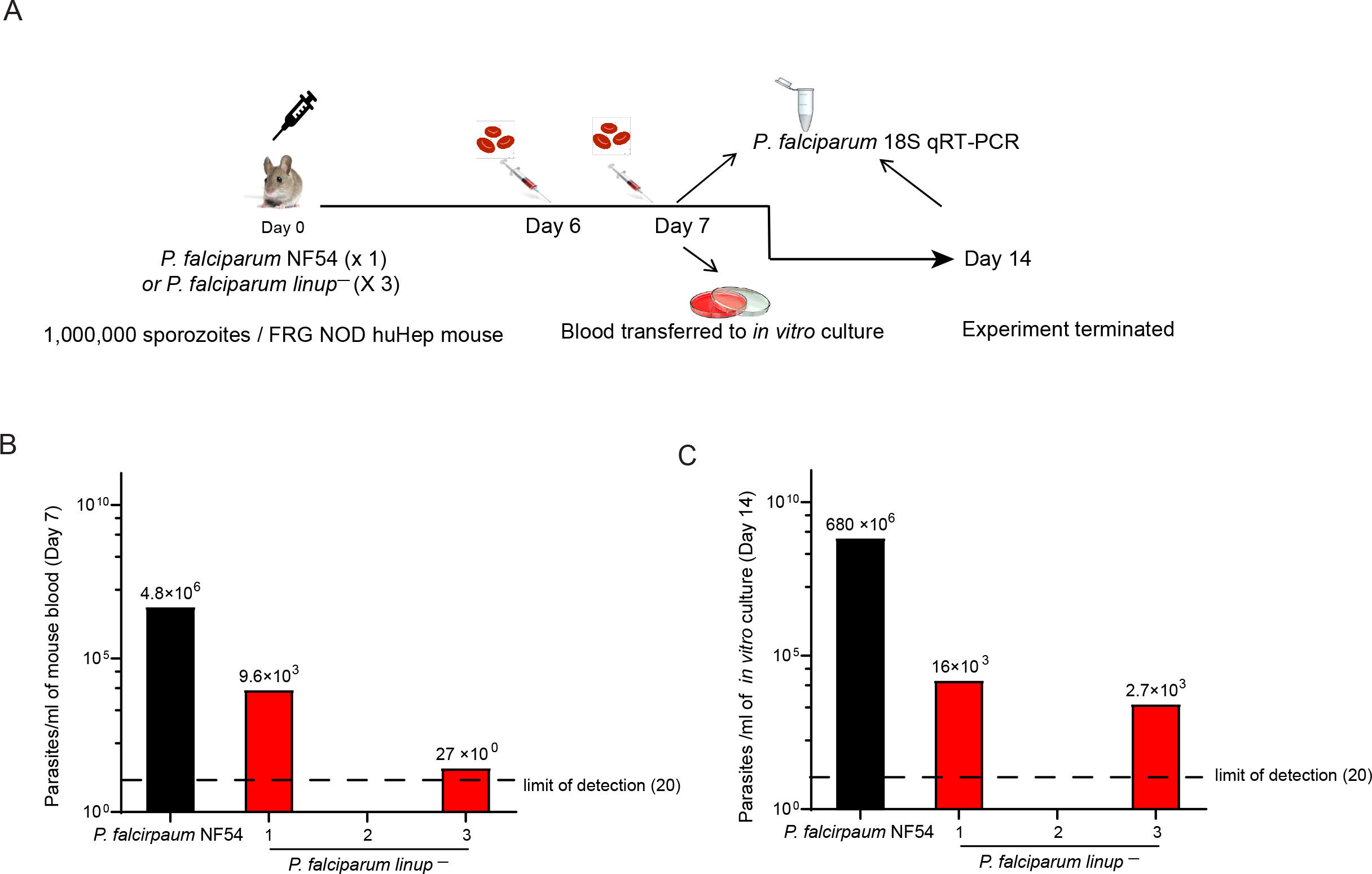
Pf *linup^−^* liver stages fail to transition to blood stages in FRG NOD huHep mice infused with human red blood cells. A. Schematic shows the experimental design of liver stage-to-blood stage transition experiments in FRG NOD huHep mice. One million sporozoites from Pf NF54 and Pf *linup^−^* were injected into one and three FRG NOD huHep mice, respectively. Mice were repopulated with human red blood cells as depicted. All mice were euthanized on day seven and 50 μl blood samples were collected from all mice for parasite load based on 18S rRNA qRT-PCR sampling. The remaining blood was transferred to in vitro culture and grown in culture for seven days. Further 50 μl blood samples were collected from in vitro culture samples for analysis of parasite 18S rRNA qRT-PCR samples seven days after transition to in vitro culture (14 days after sporozoite infection). B. Analysis of parasite load by 18 S rRNA qRT-PCR was carried out on extracted RNA from the blood of mice infected with Pf NF54 (black bar) and Pf *linup^−^* (red bars) sporozoites on day seven. There was a severe defect in the Pf *linup^−^* transition; one of three mice infected with Pf *linup^−^* sporozoites did not become patent, while the other two infected mice had an approximate 10,000-fold reduction in parasite load as compared to wildtype. C. Blood was transitioned to *in vitro* culture and parasite load was measured again after a further seven days, 14 days after sporozoite injection. Wildtype NF54 blood stages significantly replicated during the intervening seven days (black bar) whilst no significant replication was seen for Pf *linup^−^* blood stages (red bars) suggesting that the liver stage-to-blood stage transition did not lead to viable blood stage parasites.

## DISCUSSION

Genetic attenuation of Pf malaria parasites leading to liver stage arrest forms the basis of a vaccination platform that provides protection from parasite challenge ^25, 26, 42^ but the discovery of genes for specific liver stage attenuation has been difficult. The present study uncovers a novel gene deletion in both the rodent malaria parasite Py and the human malaria parasite Pf, that causes the creation of a LARC GAP. Deletion of *LINUP* in both parasites species only affected a single stage of the life cycle - the formation of exo-erythrocytic merozoites at the culmination of liver stage development. In both cases, LINUP deletion led to late liver stage and at this late stage, the liver stage parasite was reduced in size when compared to the wildtype, with no apparent formation of exo-erythrocytic merozoites.

Nevertheless, the Py *linup^−^* liver stages could transition to a patent blood stage parasitemia, demonstrating an incomplete attenuation in a subset of liver stage parasites. *LINUP*was discovered by analyzing a mature Pv liver stage transcriptome and encouragingly, top hits included genes that are known to be critical for complete liver stage development in both rodent and human malarias. We used strict criteria in selecting LINUP for this study, since we required that the gene expressed by the Pv liver stage have both Py and Pf orthologs. Of note, top hits also included PF3D7_0830300 and PF3D7_1401700 which do not have Py orthologs.

Therefore, deletion these genes from the Pf genome should be attempted, and if successful, the gene deletion phenotypes should be further studied.

Py LINUP is expressed in the nucleus from mid- to late-liver stage and its expression profile partially overlapped with the active histone 3 marker, acetylate lysine 9 (H3K9Ac) ^63^, suggesting that LINUP plays a specific role in the transcription of genes required for mid-to-late liver stage development, a time point when the liver stage parasite is undergoing both dramatic growth and differentiation into individual exoerythrocytic merozoites. We thus hypothesize that LINUP is plays a role in coordinating the transcriptional events leading to merozoite formation. Indeed, Py *linup^−^* liver stages show perturbation of merozoite formation. In addition, Py *linup^−^* liver stages also showed a decrease in size at 36- and 48-hours of development suggesting that LINUP is also involved in liver stage parasite growth. Interestingly, Pf *linup^−^* liver stages did not show a decrease in growth at five days of development (a mid-point for Pf liver stage growth) but did show a decrease liver stage growth at seven days of development, pointing to a similarity between the knockout phenotype when comparing Py with Pf. Pf *linup^−^* liver stages were also aberrant at day seven of development and like their Py counterpart showed incomplete merozoite formation and organelle division, suggesting the phenotype of the gene knockout is shared between the two species. In addition, we were unable to detect a productive Pf *linup^−^* liver stage-to-blood stage transition, suggesting the Pf LINUP, like Py LINUP, plays an important role for this transition.

Why is LINUP only required for liver stage replication but not asexual blood stage replication or sporozoite production, further stages of the life cycle that rely on rapid replication? We posit that liver stage replication, where tens of thousands of exoerythrocytic merozoites are formed, rather than 10’s of erythrocytic merozoites and 1000’s of sporozoites, requires a more coordinated control of both transcription and translation and that this requires a unique subset of genes. Interestingly, deletion of Py and Pf *PlasMei2*, an additional gene expressed exclusively during mid- to late-liver stage development also leads to a severe attenuation ^26, 45^. PlasMei2 contains a putative RNA binding domain, and its expression is associated with a granular cytoplasmic location and was hypothesized to be involved in RNA stability during liver stage maturation. It is possible that LINUP and PlasMei2 operate to regulate gene transcription and translation respectively during exo-erythrocytic merozoite formation.

Rodent malaria LARC GAP are powerful immunogens that can protect from a wildtype sporozoite challenge ^10, 43, 44^. Similarly, we have shown that Py *linup^−^* is a LARC GAP since Py *linup^−^* sporozoites infect BALB/cJ mice, develop to late liver stage and arrest. Further, a prime boost/regimen of 10,000 Py *linup^−^* sporozoites and a 10,000 wildtype sporozoite challenge gave complete protection, as has been seen for other Py LARC GAPs ^43^, suggesting that Py *linup^−^* is as effective as previously described Py LARC GAP. This thus led us to create Pf *linup^−^* and the phenotype of this GAP also suggested severe arrest late in liver stage development, demonstrating that Pf *linup^−^* is a LARC GAP worth for further pre-clinical studies Clinical studies using Pf GAP initially focused on EARD GAP, partially due to the lack of knowledge of the Pf liver stage transcriptome and proteome. These studies led to the creation of EARD GAP that have entered clinical trials and include Pf GAP3KO ^40, 41^, which has a deletion in three genes essential for early liver stage development and Pf GA1, which has deletions in two genes ^42, 64^. Pf GAP3KO and Pf GA1 have entered clinical trials. Pf GAP3KO sporozoites were administered by mosquito bite and the wildtype challenge was also administered by mosquito bite. Conversely, Pf GA1 sporozoites were isolated and cryopreserved from aseptic mosquitoes and administered intravenously, as was the wildtype sporozoite challenge. These studies demonstrated that GAP immunization could lead to sterile protection after challenge, showing the efficacy of the GAP platform ^25, 42^. However, as mentioned above, studies in rodents show that LARC GAP are superior immunogens to EARD GAP ^43^ and thus the creation of Pf LARC GAP is necessary to confirm these findings in human trials. To this end, a previous study has shown that deletion of Pf *PlasMei2* results in a severe late liver stage attenuation with no evidence of productive liver stage-to-blood stage transition using a FRG NOD huHep mouse model ^26^ and thus Pf *PlasMei2*is a LARC GAP. However, analysis of Py *plasmei2^−^*, which also arrests late in liver stage development showed that at very high doses, mice became patent after infection ^10^. Thus, there is a concern that Pf *plasmei2^−^* may breakthrough to patent parasitemia if this LARC GAP is not completely attenuated. Based on these findings, the creation of a safe and immune protective Pf LARC GAP may rely on the deletion of a second gene from the more Pf *plasmei2^−^* LARC GAP to ensure complete attenuation. The results from this study suggest that the deletion of Pf *LINUP* from the Pf *plasmei2^−^* LARC GAP could further attenuate this LARC GAP, creating completely attenuated parasite for further pre-clinical and clinical development. To aid in the discovery of further Pf gene deletions for the creation of Pf LARC GAP, a comprehensive Pf liver stage transcriptome would be a vital resource for revealing novel candidates for further study.

## MATERIALS AND METHODS

### Study Approval

This study was carried out at the Seattle Children’s Research Institute in accordance with the recommendations of the NIH Office of Laboratory Animal Welfare standards (OLAW welfare assurance #D16-00119). The Seattle Children’s Research Institute Institutional Animal Care and Use Committee (IACUC) reviewed all protocols utilizing research animals used in this study. The IACUC protocols used in this study were AUCU00505 (SW, and BALB/c mice) and ACUC00480 (FRG NOD huHep mice).

### Experimental animals

Six- to eight-week-old female Swiss Webster (SW) mice were purchased from Envigo and used for Py parasite life cycle maintenance and production of transgenic parasites. Six- to eight-week-old female BALB/cJ and BALB/cByJ mice were purchased from Jackson Laboratories and used for assessments of parasite infectivity, indirect immunofluorescence assays (IFA), as well as the attenuation and ability of Py *linup*^-^ parasite to act as an experimental vaccine. For studies of Pf *linup*^-^, FRG NOD huHep mice (female and >4 months of age) were purchased from Yecuris, Inc. Repopulation of human hepatocytes was confirmed by measuring human serum albumin levels, and only animals with human serum albumin levels >4 mg/mL were used, corresponding to >70% human hepatocyte repopulation. FRG NOD huHep mice were maintained on drinking water containing 3% Dextrose and were cycled on 8 mg/L NTBC once a month for 4 days to maintain hepatocyte chimerism. Py 17XNL wildtype and transgenic parasites were cycled between SW mice and *Anopheles stephensi* mosquitoes for the purposes of sporozoite production. Infected mosquitoes were maintained on sugar water at 24°C and 70% humidity.

### Creation of a Py LINUP^mCherry^ *and* Py *linup*^-^ parasites

All oligonucleotide primers used for the creation and analysis of Py LINUP^mCherry^ and Py *linup*^-^ are detailed in **Table S1**. Creation of Py LINUP^mCherry^ utilized double crossover homologous recombination using modified plasmid pL0005 (obtained through the MR4 as part of the BEI Resources Repository, NIAID, NIH: *Plasmodium berghei* pL0005, MRA-774, deposited by AP Waters), which allowed for the addition of a mCherry epitope tag to the carboxy terminus of LINUP. The resultant Py LINUP^mCherry^ parasite expresses a single copy of LINUP with an mCherry tag under the control of its endogenous promoter. Two individual Py LINUP^mCherry^ clones underwent life cycle phenotypic analysis.

Deletion of Py *LINUP* (PlasmoDB identifier PY17X_1465200) was achieved using CRISPR/Cas9 technology ^65^. In brief, *LINUP* was deleted using double crossover homologous recombination and complementary regions of *LINUP*upstream and downstream of the open reading frame were ligated into plasmid pYC L2 to create pYC_LINUP. pYC_LINUP was transfected into the blood stage schizonts of Py 17XNL. After transfection and intravenous injection into SW mice, pyrimethamine was used for the positive selection and downstream cloning of recombinant parasites using standard techniques ^57^. Gene knockout was confirmed by PCR using methodology we have used on multiple occasions (see ^66^ for a recent example). This led to the creation of the Py *linup*^-^ and two separate knockout clones from two independent transfections were initially phenotypically analyzed throughout the life cycle.

Phenotypic analysis of blood stage Py *linup^−^*. To assay the growth of Py 17XNL and Py *linup^−^* blood stages, blood was removed from infected SW mice when parasitemia was between 0.5 and 1.5%. Blood was diluted in RPMI 1640 medium (HyClone, Logan, UT) so that 100 μl contained 10^6^ parasites. SW mice (five in each group) were then injected intravenously with the infected red blood cells, and the percent parasitemia was monitored every other day until day 11.

### Sporozoite inoculation and challenge

Py *linup*^-^ and Pf *linup*^-^ sporozoites were isolated from the salivary glands of infected *A. stephensi* mosquitoes between 14 and 18 days after the infectious blood meal and injected intravenously into the tail vein of recipient mice. For assessment of Py *linup*^-^ attenuation, sporozoites were injected intravenously into highly susceptible BALB/cByJ mice ^58^. Liver stage-to-blood stage transition (blood stage patency) was assessed by Giemsa-stained thin blood smear starting at day three after inoculation and ending at day 14, at which time, a negative smear was attributed to complete attenuation. For assessment of Pf *linup*^-^ attenuation, sporozoites were injected intravenously into FRG huHep mice and human red blood cells were intravenously injected on the sixth and seventh day after infection.

For immunizations, SW, BALB/cJ and BALB/cByJ mice were primed and boosted with Py sporozoites and subsequently challenged with 10,000 wildtype Py XNL sporozoites.

Breakthrough to blood stage patency was assessed by Giemsa-stained thin blood smear starting at day three after challenge and ending at day 14, at which time, a negative smear was attributed to complete protection. Mice immunized only with mosquito salivary gland extract were used as controls.

#### *In vitro* culturing of Pf parasite lines

Pf lines and were cultured in custom-made RPMI 1640 media containing hypoxanthine, sodium bicarbonate, and 4-(2-hydroxyethyl)-1- piperazineethanesulfonic acid (Invitrogen, Life Technologies, Grand Island, NY) supplemented with 5% human serum (Valley Biomedical), 5% AlbuMAX (Invitrogen, Life Technologies, Grand Island, NY) and gentamycin (Invitrogen), in presence of O^+^ blood (Valley Biomedical) at 3 – 5% hematocrit, in an atmosphere of 5% CO2, 5% O2, and 90% N2.

#### Creation of the Pf *linup*^-^ parasite

The CRISPR-Cas9 vector for the Pf gene targeting strategy have been previously described ^26, 67^. The pFC-LINUP plasmid was generated by cloning 5’ and 3’ homology arms flanking the Pf *LINUP* gene (PF3D7_1249700) into the pFC vector followed by cloning the guide RNA sequence. 5% sorbitol synchronized ring stage Pf NF54 parasites at 5-8% parasitemia were electroporated with 100 µg of pFC-LINUP plasmid at 0.31 kV and 950 µF using a BioRad Gene Pulser (BioRad, Hercules, CA). Positive selection with 8 nM WR99210 (WR; Jacobus Pharmaceuticals, Princeton, NJ) was administered 24 hours after the transfection and kept for five days. Drug resistant parasites were rescued after 21 days. These parasites were screened by PCR using gene specific primers (Table S1) to confirm gene deletion.

#### Pf parasite cloning by limiting dilution

Pf *linup^−^* parasites were cloned out of the mixed parasite pool by limiting dilution in 96-well flat bottomed plates (4). Cultures were serially diluted and then plated at a density of 0.3 parasites per well in a 100 µl volume at 2% hematocrit. Cultures were fed once a week and fresh blood was added every three – four days at 0.3% hematocrit. The wells were screened for presence of parasites using primers to 18S RNA on day (Table S1) [18SF (5’ AACCTGGTTGATCCAGTAGTCATATG 3’) and 18SR (5’CCAAAAATTGGCCTTGCATTGTTAT 3’)]. Positive wells were expanded and screened for recombination using the PCR strategies used above. Pf *linup^−^* clones B1 and B4 were used for further phenotypic analysis.

#### Gametocyte culture and mosquito infections

Pf wildtype NF54 and Pf *linup^−^* cultures were grown in gametocyte media containing RPMI 1640 media containing hypoxanthine, sodium bicarbonate, and 4-(2-hydroxyethyl)-1-piperazineethanesulfonic acid (Invitrogen, Life Technologies, Grand Island, NY) supplemented with 10% human serum (Valley Biomedical) in presence of O+ blood (Valley Biomedical) at 4% hematocrit, in an atmosphere of 5% CO2, 5% O2, and 90% N2. Gametocyte cultures were initiated at 4% hematocrit and 0.8%–1% parasitemia (mixed stages) and maintained for up to 15 days with daily medium changes. On days 15 – 17 post set up, stage V gametocytemia was evaluated by Giemsa-stained thin blood smears. Mature gametocyte cultures were spun down at 800 *g* for 2 mins at 37°C to pellet infected red blood cells. A volume of pre-warmed 100% human serum equal to the packed red blood cell volume was added to resuspend the red blood cell pellet. The resuspended pellet was diluted to a gametocytemia of 0.4 – 0.5% with pre-warmed feeding media containing 50% human serum and 50% human red blood cell and immediately fed to non–blood fed adult female mosquitoes 3 to 7 days after hatching by membrane feeding assay. Mosquitoes were allowed to feed through Parafilm for up to 20 minutes. Following blood feeding, mosquitoes were maintained for up to 19 days at 27°C, 75% humidity, and provided with 8% dextrose solution in PABA water. Oocyst prevalence was checked on day 7 – 9 post feed by dissecting approximately 12 midguts per cage. Sporozoite numbers were detected by dissecting and grinding salivary glands in Schneider′s Insect Medium (Sigma) on days 14 – 18 post feed.

**Phenotypic analysis of blood stage Pf *linup^−^***

Ring stage parasites of Pf NF54 WT and Pf *linup^−^* were set up at 0.5% parasitemia and 3% hematocrit in media containing 5% Albumax and 5% human serum. Fresh media was repleted daily. Growth was monitored over three replication cycles by making thin blood smeared every other day. Cultures were cut back by 3 – 5 folds after the second replication cycle to prevent parasite cultures from crashing. The parasitemia after the third replication cycle was calculated by multiplying the parasitemia based on blood smear with the diluted fold change.

#### Phenotypic analysis of Pf *linup^−^* parasites in FRG NOD huHep mice

Pf NF54 and Pf *linup^−^* sporozoites were isolated from salivary glands of infected *Anopheles stephensi* mosquitoes. For analysis of Pf *linup^−^* liver stage, 1 x 10^6^ Pf NF54 and 1 x 10^6^ Pf *linup^−^* sporozoites were injected intravenously (retro-orbital) into four FRG NOD huHep mice per group. Livers were harvested on days 5 and 7 and used for IFA. To evaluate for blood stage transition of Pf *linup^−^* in FRG NOD huHep mice, 1 x 10^6^ Pf NF54 and Pf *linup^−^* sporozoites from clone were injected intravenously into one and three FRG NOD huHep mice per group respectively. On day 6 and 7, 400 µl of 70% RBCs were injected intravenously to enable transition of liver stage parasites to blood. Four hours after human RBC repopulation on day 7, mice were euthanized, blood was collected by cardiac puncture and 50µl of blood from each mouse was used for qRT-PCR analysis to detect parasite 18S RNA. Blood was washed three times in asexual media, a volume of human RBCs equal to the packed RBC volume was added and blood was transferred to in vitro culture. Fresh media was replaced daily, and cultures were analyzed every 2-3 days by thick smear for presence of parasites for up to 7 days (14 days after sporozoite inoculation).

Samples from in vitro culture were analyzed for presence of 18S rRNA by qRT-PCR after 7 days.

#### *Plasmodium* 18S rRNA qRT-PCR quantification of parasite load

For quantification of *Plasmodium* 18S rRNA from mouse blood, 50 µL of whole blood was added to 2 mL of NucliSENS lysis buffer (bioMérieux, Marcy-l’Étoile, France) and frozen immediately at -80°C. One mL of lysate was subsequently processed using the Abbott m2000sp using mSample RNA preparation kit (Abbott, Niles, IL). The eluate was tested by qRT-PCR for the pan-*Plasmodium* 18S rRNA target. The qRT-PCR reaction was performed using 35 µL SensiFAST™ Probe Lo- ROX One-Step Kit (Bioline, Taunton, MA) and 15 µL of extracted eluate. Plasmodium 18S rRNA primers/probes (LCG BioSearch Technologies, Novato, CA) were as follows: Forward primer PanDDT1043F19 (0.2 µM): 5’-AAAGTTAAGGGAGTGAAGA-3’; Reverse primer PanDDT1197R22 (0.2 µM): 5’-AAGACTTTGATTTCTCATAAGG-3’; Probe (0.1 µM): 5’-[CAL Fluor Orange 560]- ACCGTCGTAATCTTAACCATAAACTA[T(Black Hole Quencher- 1)]GCCGACTAG-3’[Spacer C3]). Cycling conditions were RT (10 min) at 48°C, denaturation (2 min) at 95°C and 45 PCR cycles of 95°C (5 sec) and 50°C (35 sec).

### Immunofluorescence assay (IFA)

For studies of Py LINUP, BALB/cByJ mice were injected intravenously with approximately 0.5 x 10^6^ Py LINUP^mCherry^ and sporozoites Py *linup*^-^ sporozoites and livers were harvested from euthanized mice at time points post infection. For studies of Pf *linup*^-^, human liver chimeric FRG huHep were injected intravenously with approximately 0.5 x 10^6^ sporozoites and livers were harvested from euthanized mice at time points post infection. For both Py and Pf studies, wildtype parasite infections were used as controls.

Livers were perfused with 1×PBS, fixed in 4% v/v paraformaldehyde (PFA) in 1×PBS and lobes were cut into 50 μm sections using a Vibratome apparatus (Ted Pella, Redding, CA). For IFA, sections were permeabilized in 1×TBS containing 3% v/v H_2_O_2_ and 0.25% v/v Triton X-100 for 30 min at room temperature. Sections were then blocked in 1×TBS containing 5% v/v dried milk (TBS-M) for at least 1 h and incubated with primary antibody in TBS-M at 4°C overnight. After washing in 1×TBS, fluorescent secondary antibodies were added in TBS-M for 2 h at room temperature in a similar manner as above. After further washing, the section was incubated in 0.06% w/v KMnO_4_ for 2 min to quench background fluorescence. Sections were then washed with 1×TBS and stained with 1 μg/ml 4′, 6-diamidino-2-phenylindole (DAPI) in 1× TBS for 5–10 min at room temperature to visualize DNA and mounted with FluoroGuard anti-fade reagent (Bio-Rad, Hercules, CA).

Primary antibodies used included Pf CSP clone 2A10 (1:500), Py BiP polyclonal (1:1000), Py ACP (1:500), *P. vivax* mHSP70 (1:1000), *P. vivax* mHSP60 (1:1000) and Pf MSP-1 clone 12.10 (1:500, European Malaria Reagent Repository), mCherry clone 16D7 (1:500, Thermo Scientific), Py mTIP rabbit polyclonal (1:500) and histone H3K9ac (acetylated lysine 9of histone 3) antibody (Genetex # GTX630554, 1:250). Secondary antibodies include donkey anti-mouse 488, donkey anti-mouse 594, donkey anti-mouse 647, donkey anti-rabbit 488, donkey anti-rabbit 594, donkey anti-rabbit 647 and donkey anti-rat 594. All secondary antibodies were used at a dilution of 1:1000.

#### Microscopy

All images were acquired using the Stellaris 8 confocal microscope (Leica Microsystems) with 63x water objective and processed using the Lightning software (Leica Microsystems). For quantification of parasite liver stage size, the parasite was assumed to be elliptical in shape and therefore area was calculated from its longest (a) and shortest (b) circumferential diameter (πab).

#### Quantification and statistical analysis

Where appropriate, quantification was represented by the mean ± standard deviation. Calculations and statistical tests indicated in the figure legends were performed using GraphPad Prism Software. The statistical tests used was two-way ANOVA. *P<0.05, ** P<0.01, ***P<0.001. A P value>0.05 was considered not significant.

## FUNDING INFORMATION

The work was funded by NIH/NIAID grants R01_AI125706 and U01_ AI155335 awarded to SHIK.

### ACKOWLEDGEMENTS

We would like to thank Seattle Children’s Research Institute insectary employees for their help in maintaining parasites in the mosquito vector. We would also like to thank Seattle Children’s Research Institute vivarium staff for their help with rodent maintenance. SHIK, AMV and DG conceived the experimental plan. AMV, DG, SA, KO, WB, and JA carried out experiments.

AMV, SHIK and DG wrote the paper.

---

**Table S1.**
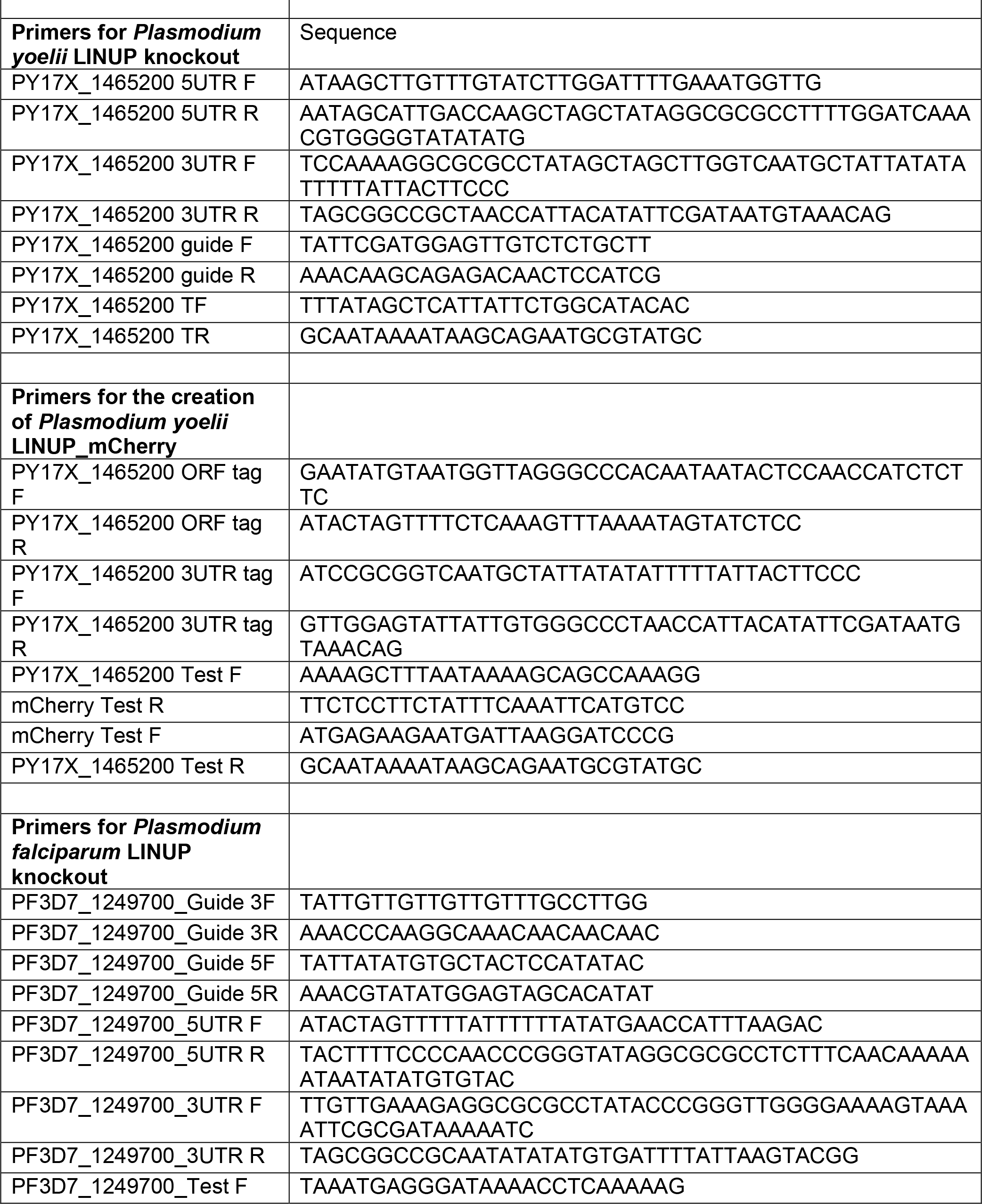
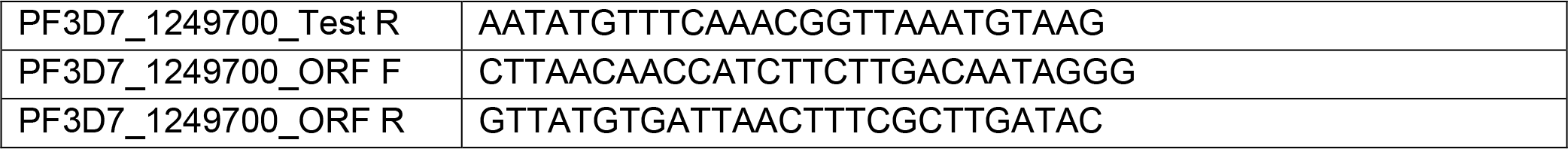
Oligonucleotide primers used in the study.

