## Supplemental Figure and text for "A conserved *Plasmodium* protein that localizes to liver stage nuclei is critical for late liver stage development"

Supplementary figure 1

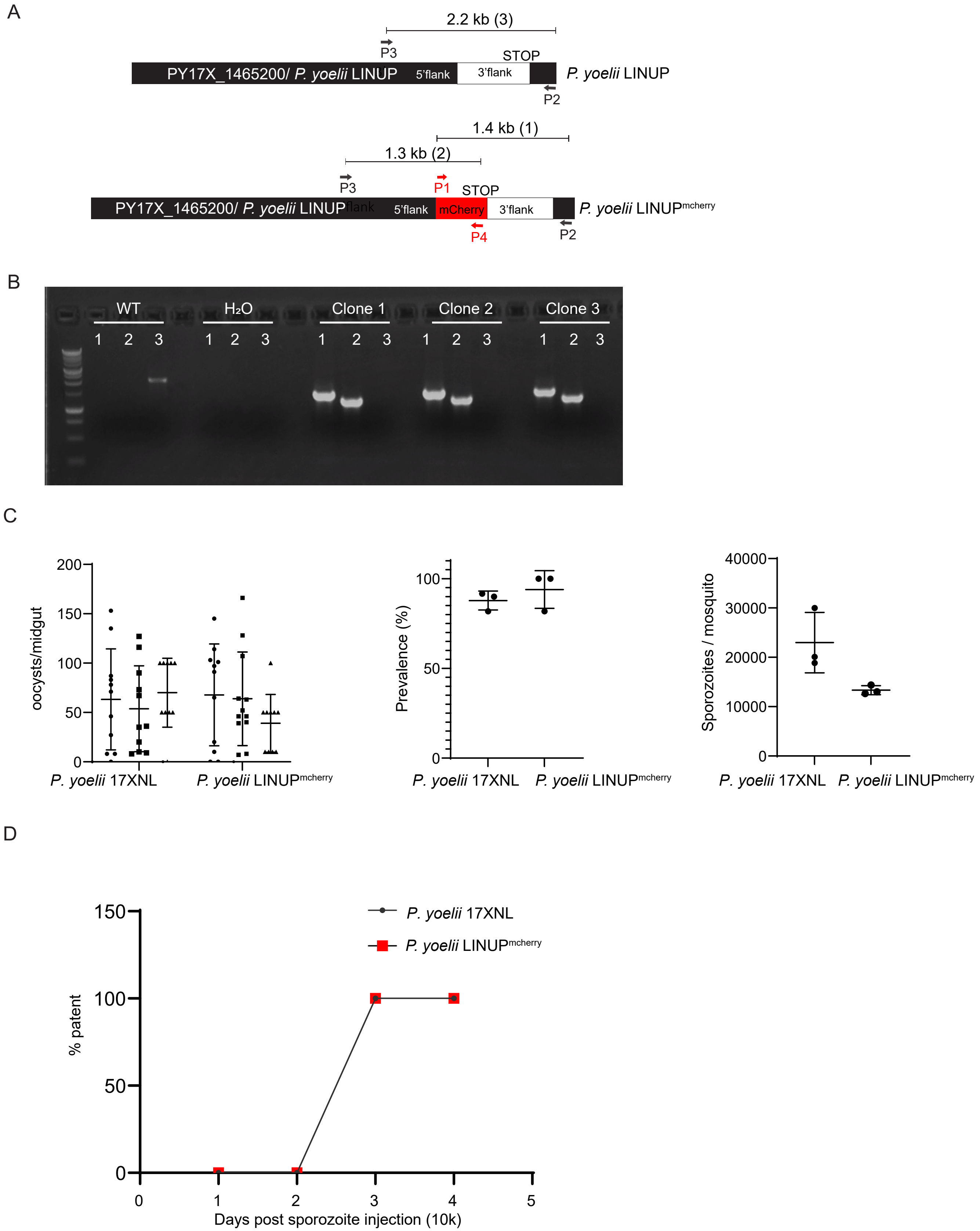

**Supplementary Figure 1. Py LINUP$^{mCherry}$ parasites undergo normal development throughout the life cycle.** A. The schematic depicts the generation of PY17X_1465200$^{mCherry}$ (Py LINUP$^{mCherry}$) parasites using CRISPR/Cas9-mediated gene editing. Primers used to verify the insertion of the mCherry tag are indicated and the sizes of the PCR amplicons are shown in kilobases. B. Agarose gel electrophoresis shows the PCR products 1, 2 and 3 corresponding to the insertion of mCherry tag to the C-terminus of PY17X_1465200 in three independent clones. C. Py LINUP$^{mCherry}$ did not have defects in sporogenesis as counts for oocysts/midgut, oocyst prevalence and salivary gland sporozoites/mosquito were comparable to Py wildtype. Data is represented as mean +/- SD, *n*=3 biological replicates. Statistical analysis was carried out using two-way ANOVA using Tukey's multiple comparison test. P>0.05 is taken as not significant. D. Py LINUP$^{mCherry}$ did not have any defects in liver stage development since SW mice challenged with 10,000 Py LINUP$^{mCherry}$ sporozoites successfully transitioned from the liver stage-to-blood stage of infection and all five mice were patent on day 3, as did mice infected with Py wildtype.
